# Acute Stress Desensitizes Hypothalamic CRH Neurons to Norepinephrine and Physiological Stress

**DOI:** 10.1101/2019.12.31.891408

**Authors:** Zhiying Jiang, Chun Chen, Grant L. Weiss, Xin Fu, Marc O. Fisher, John C. Begley, Carly R. Stevens, Laura M. Harrison, Jeffrey G. Tasker

## Abstract

Noradrenergic afferents to corticotropin releasing hormone (CRH) neurons of the hypothalamic paraventricular nucleus (PVN) provide a major excitatory drive to the hypothalamic-pituitary-adrenal (HPA) axis via α1 adrenoreceptor activation. The ascending noradrenergic afferents are recruited preferentially by physiological, rather than psychological, stress modalities.

Glucocorticoids secreted in response to HPA activation feed back onto the hypothalamus to negatively regulate the HPA axis, providing a critical autoregulatory constraint that prevents glucocorticoid overexposure. Whether differential negative feedback mechanisms target stress modality-specific HPA activation is not known. Here, we reveal a desensitization of the α1 adrenoreceptor activation of the HPA axis following acute stress that is mediated by rapid glucocorticoid regulation of adrenoreceptor trafficking. Prior stress desensitized the HPA axis to subsequent physiological, but not psychological, stress. Our findings demonstrate rapid glucocorticoid suppression of adrenoreceptor signaling in CRH neurons that is specific to physiological stress activation, and reveal, therefore, a rapid, modality-selective glucocorticoid feedback mechanism.

## INTRODUCTION

Activation of the hypothalamic-pituitary-adrenal (HPA) axis is part of Selye’s generalized stress response described over 50 years ago^1^. How this response that stimulates adrenal secretion of glucocorticoids discriminates between different stress modalities is not known. Here, we targeted the noradrenergic excitatory drive to the HPA axis, which is activated preferentially by systemic, physiological stress stimulation^2-4^. The PVN corticotropin releasing hormone (CRH) neurons, the primary effector cells of the HPA axis, are innervated by noradrenergic afferents from the A2 cell group of the nucleus of the solitary tract (NTS), the activation of which robustly stimulates the HPA axis^5-7^. Multiple studies have suggested that this afferent pathway provides the main excitatory drive to the HPA axis during systemic stress exposure, such as that triggered by glycemic or immune challenge^3,4,8,9^. We recently characterized a non-canonical α1 adrenoreceptor-mediated noradrenergic activation of the corticotropin releasing hormone (CRH) neurons of the hypothalamic paraventricular nucleus (PVN) that is mediated by the activation of local glutamate circuits following postsynaptic α1 receptor activation, dendritic peptide release, and retrograde glial-neuronal signaling^10^. We also provided evidence for the monosynaptic co-release of glutamate with norepinephrine from the noradrenergic afferents^10^. Thus, the noradrenergic afferents stimulate PVN CRH neurons by direct monosynaptic glutamate co-release and via α1 receptor-mediated recruitment of local glutamatergic circuits.

Glucocorticoid negative feedback inhibits the activation of the HPA axis^11^, which protects the organism from the damaging effects of sustained corticosteroid exposure. Previous studies showed that elevated plasma corticosterone decreases the PVN neuronal response to NE and HPA axis activation. Systemic pretreatment with the synthetic glucocorticoid dexamethasone inhibited the ACTH and corticosterone responses to noradrenergic afferent stimulation in a dose-dependent manner^12^. Manipulating corticosterone levels by adrenalectomy, corticosterone pellet supplementation, and acute stress demonstrated that plasma corticosterone levels are inversely correlated with PVN α1b adrenoreceptor mRNA expression^13^. Norepinephrine excites a subset of PVN parvocellular neurons, which is blocked by cortisol pretreatment^14^. In addition, adrenalectomy increased the α1 adrenoreceptor-mediated excitation of PVN parvocellular neurons, which was reversed by corticosterone replacement^15^. These findings together suggest an inhibitory effect of glucocorticoids on the NE-induced excitation of hypothalamic PVN neurons, and that the NE-induced excitation of the HPA axis activity is highly sensitive to corticosteroid levels, suggesting it is a target of glucocorticoid negative feedback.

The mechanisms of glucocorticoid suppression of NE signaling to the HPA axis represent a critical pharmacotherapeutic target because of the association of HPA dysfunction with stress disorders, however these mechanisms are not known. We investigated whether stress-induced glucocorticoids inhibit the NE-induced excitation of CRH neurons and HPA activation using patch-clamp recordings in acute mouse brain slices, live-cell imaging in a hypothalamic cell line, and *in vivo* physiological analyses. Surprisingly, we found that stress desensitizes PVN CRH neurons to NE via glucocorticoid regulation of α_1_-adrenoceptor trafficking. The stress/glucocorticoid-induced NE desensitization of the CRH neurons causes a suppression of HPA activation by physiological stress, but not by psychological stress, revealing a stress modality-selective glucocorticoid negative regulation of the HPA axis.

## RESULTS

### Stress-state dependence of the NE α1 adrenoreceptor response in CRH neurons

Norepinephrine elicits a concentration-dependent excitation of PVN CRH neurons by stimulating local glutamate circuits via activation of postsynaptic α_1_ adrenoreceptors in the CRH neurons leading to the retrograde activation of a glial-neuronal circuit^10^ (Fig. S1A). Here, we tested for the stress-state dependence of the NE response in PVN CRH neurons by subjecting mice to an acute stress before performing whole-cell recordings in acute slices. In PVN CRH neurons from control mice, bath application of NE (100 μM) caused a robust increase in the frequency of spontaneous EPSCs (sEPSCs) (335.2 ± 61.5% of baseline, p < 0.001, n = 17) (Fig. 1A-C), but had no effect on the sEPSC amplitude or decay^10^. In CRH neurons from mice subjected to a 30-min restraint stress, the NE response was suppressed, resulting in a non-significant increase in sEPSC frequency (145.9 ± 39.4% of baseline, p = 0.29, n = 7) (Fig. 1A-C). No difference was detected between CRH neurons from unstressed mice and stressed mice in the basal sEPSC frequency (unstressed: 1.56 ± 0.10 Hz, n = 47; stressed: 1.36 ± 0.16 Hz, n = 12, p = 0.46), amplitude (unstressed: 19.12 ± 0.50 pA, n = 47; stressed: 17.64 ± 1.03 pA, n = 12, p = 0.13), or decay time (unstressed: 2.18 ± 0.06 ms, n = 47; stressed: 2.05 ± 0.12 ms, n = 12, p = 0.38).

**Figure. 1.**
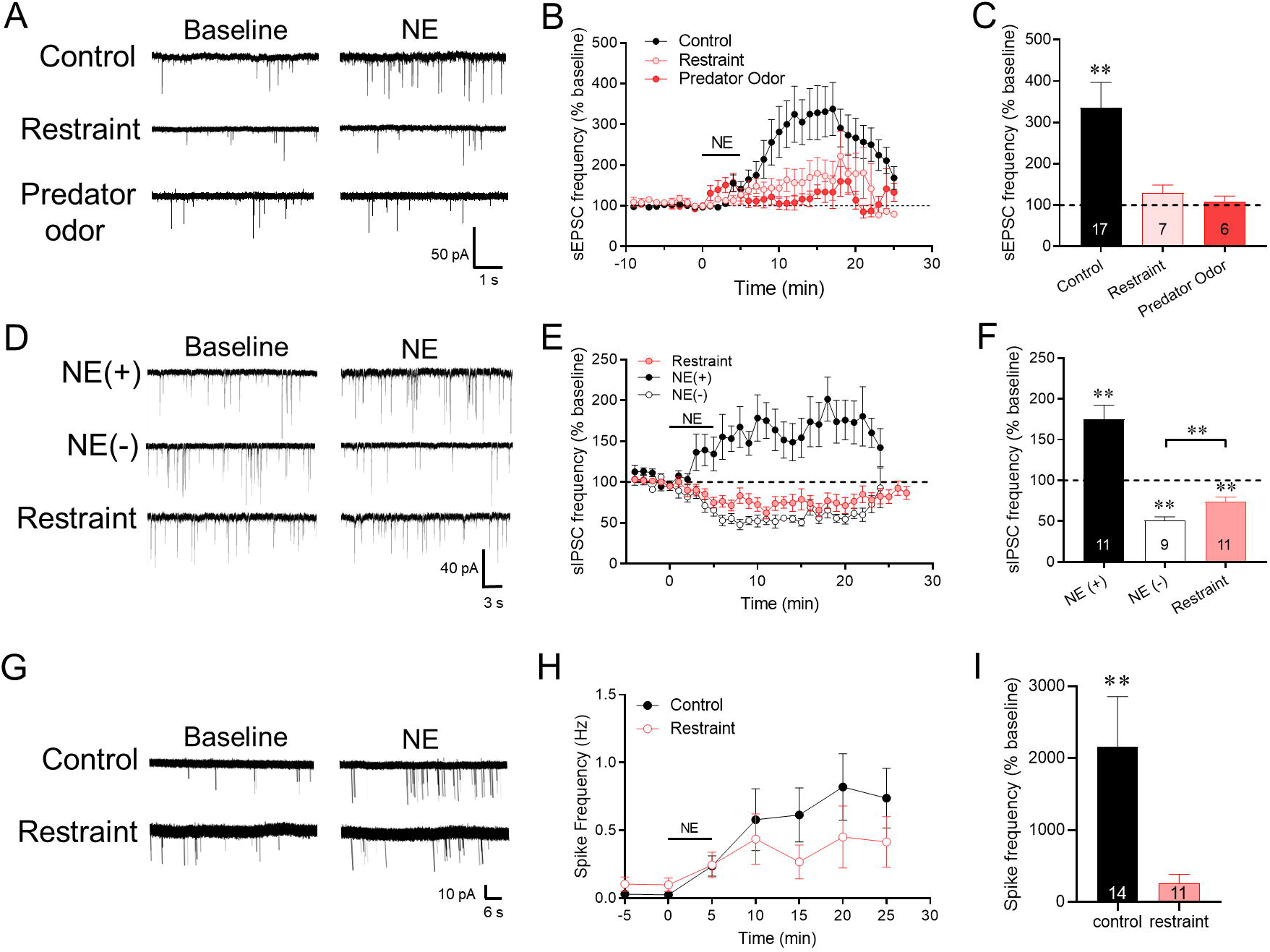
The NE-induced facilitation of glutamate release is stress-sensitive. **A**. Representative traces of the NE-induced increase in sEPSC frequency recorded in CRH neurons from unstressed mice and mice exposed to 30-min restraint or 30-min predator odor. **B**. Time course of mean normalized sEPSC frequency response (% of baseline) to NE (100 μM) in CRH neurons from control mice and mice subjected to restraint or predator odor. **C**. Summary plot of mean normalized sEPSC frequency response to NE in CRH neurons from control mice and mice subjected to restraint and predator odor. **D**. Representative traces of the increase in sIPSC frequency and the decrease in sIPSC frequency seen in PVN CRH neurons in response to NE (NE(+) and NE(-), respectively), and the sIPSC response to NE in a CRH neuron from a mouse subjected to 30-min restraint stress (Restraint). NE had a dual effect on sIPSC frequency in CRH neurons from control mice, causing an increase in sIPSC frequency (NE(+)) in some CRH neurons and a decrease in sIPSC frequency (NE(-)) in others^10^. **E**. Time course of the mean normalized sIPSC frequency responses (% of baseline, 1-min bins) to NE in CRH neurons from control mice and mice subjected to a 30-min restraint stress. The NE(+) response was lost and only the NE(-) response was recorded in CRH neurons from stressed mice. **F**. Summary plot of the mean normalized sIPSC frequency responses to NE showing the dual response in CRH neurons from control mice (NE(+) and NE(-)) and the restraint stress block of the NE-induced facilitation of sIPSCs. **, p < 0.01. **G**. Representative loose-seal patch clamp recordings of the NE effects on CRH neuron spiking in slices from unstressed mice (Control) and restraint-stressed mice (Restraint). **H**. Time course of the effects of NE on mean normalized spiking frequency (5-min bins) in CRH neurons from unstressed (Control) and restraint-stressed mice (Restraint). **I**. Summary plot of the NE-induced change in mean normalized spiking frequency (% change) compared to the baseline frequency. ** p < 0.01.

To determine whether the restraint stress-induced desensitization of the CRH neurons to α1 adrenoreceptor activation extends to other types of acute stressors, we also tested a 30-min exposure to a predator odor^16^. Bobcat urine exposure prior to sacrifice completely blocked the NE-induced facilitation of sEPSC frequency in CRH neurons (108.3% ± 13.82% of baseline, p = 0.57, n = 6) (Figure 1B, C).

Norepinephrine has a dual effect on inhibitory synaptic inputs to PVN CRH neurons: 1) postsynaptic α1 receptor-mediated retrograde activation of local GABA circuits that facilitates inhibitory inputs, and 2) presynaptic α2 receptor-mediated suppression of GABA release that suppresses inhibitory inputs (Fig. S1B); approximately half of CRH neurons respond with α1 receptor-mediated retrograde facilitation of GABA inputs, while all the CRH neurons respond with presynaptic α2 receptor-dependent suppression of GABA inputs, which is masked in neurons showing a facilitation^10^. The NE activation of local inhibitory inputs to CRH neurons is also mediated by retrograde neuronal-glial signaling (Fig. S1B), so we tested for the stress desensitization of the α1 receptor-induced facilitation of spontaneous inhibitory postsynaptic currents (sIPSCs). Following restraint stress, none of the CRH neurons responded to NE (100 μM) with an increase in sIPSC frequency, all 11 cells responding with a decrease in sIPSC frequency (74.23% ± 5.24% of baseline, n = 11, p < 0.01) (Fig. 1D-F). Therefore, prior stress exposure suppressed the NE-induced facilitation of excitatory and inhibitory synaptic inputs to CRH neurons, which suggested a desensitization of postsynaptic α1 adrenoreceptors.

Norepinephrine stimulates an increase in spiking activity in CRH neurons that is driven by the NE-induced increase in excitatory synaptic inputs^10^. We tested whether the stress-induced suppression of excitatory synaptic inputs reduces the NE activation of the CRH neurons with loose-seal patch-clamp recordings. Slices were incubated in high-[K^+^] solution (10 mM) to stimulate spontaneous spiking; the mean baseline spiking frequency in CRH neurons from stressed mice (0.10 ± 0.05 Hz, n = 11) was ∼4-fold higher than in CRH neurons from control mice (0.03 ± 0.02 Hz, n = 14), but this difference did not reach statistical significance (p = 0.15). Bath application of NE (100 *µ*M, 5 min) caused a 25-fold increase in the spiking frequency in CRH neurons from control mice (from 0.03 ± 0.02 Hz to 0.74 ± 0.22 Hz, n = 14, p < 0.01) (Fig 1G-I), and a 4-fold increase in the spiking frequency in CRH neurons from restraint-stressed mice, which did not reach statistical significance (from 0.10 ± 0.05 Hz to 0.42 ± 0.19 Hz, n = 11, p = 0.067) (Fig. 1G-I).

### Glucocorticoid suppression of NE excitation of CRH neurons

Several studies have shown that acute glucocorticoid pretreatment suppresses stress-induced HPA activation^17-19^. Here, we tested whether the acute stress-induced suppression of the NE activation of excitatory synaptic inputs to PVN CRH neurons is due to increased plasma glucocorticoid levels induced by HPA activation.

The 11-β-hydroxylase inhibitor metyrapone suppresses stress-induced corticosterone synthesis (Fig. S2). As before, the NE-induced increase in sEPSC frequency was abolished in CRH neurons from I.P. vehicle-injected, restraint-stressed mice (128.5 ± 21.0%, n = 8, p = 0.22). The NE response was partially restored in CRH neurons from stressed mice pretreated with metyrapone (100 mg/kg IP) (204.2 ± 32.0% of baseline, p = 0.012, n = 9) (Fig 2A, B), suggesting that the stress-induced suppression of NE activation of excitatory synaptic inputs to the CRH neurons is dependent on stress-induced glucocorticoid secretion.

**Figure 2.**
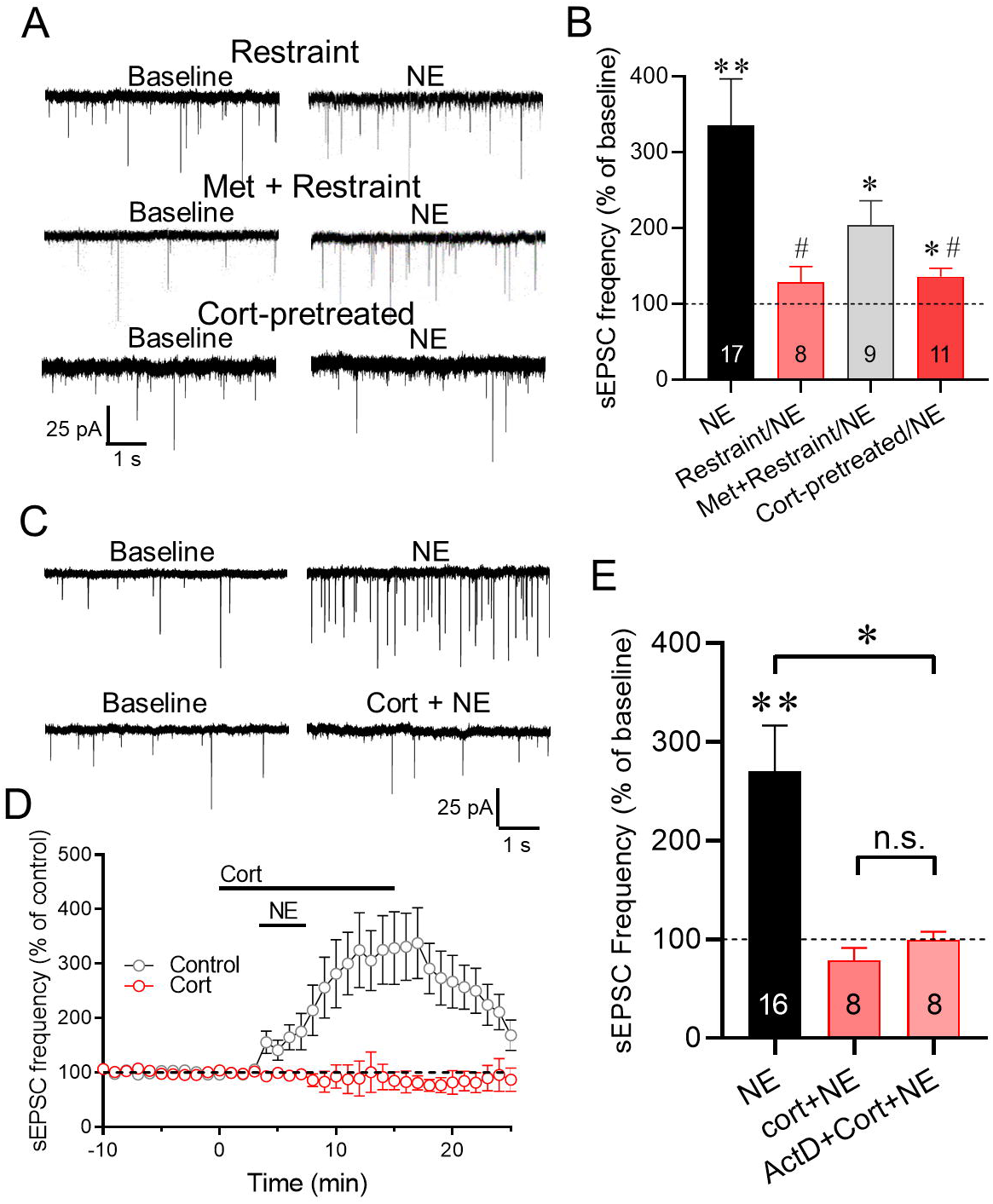
Acute stress and glucocorticoid inhibit the NE-induced increase in excitatory synaptic inputs to CRH neurons. **A**. Representative recordings of sEPSCs at baseline and following NE application in CRH neurons in slices from restraint-stressed mice after I.P. saline injection (Restraint) or metyrapone injection (Met+Restraint), and in slices from control mice that were preincubated for 5 min in corticosterone 2 h prior to NE application (Cort-pretreated). **B**. Summary plot of the NE-induced increase in mean normalized sEPSC frequency (% of baseline) in CRH neurons from unstressed control mice (NE), from mice subjected to restraint stress following I.P. saline injection (Restraint/NE), from mice subjected to restraint stress following I.P. metyrapone injection (Met+Restraint/NE), and from slices from control mice preincubated in corticosterone (Cort-pretreated/NE). The NE-induced increase in sEPSC frequency was blocked in CRH neurons following in-vivo restraint and preexposure of slices to corticosterone, and was partially rescued with the corticosterone synthesis inhibitor metyrapone. **C**. Representative recordings of the sEPSC response in CRH neurons to NE in the absence (NE) and in the presence of corticosterone (Cort+NE). **D**. Time course of the change in the mean normalized sEPSC frequency (% of baseline) in CRH neurons in response to NE alone (Control) and to corticosterone and NE (Cort). The NE-induced increase in sEPSC frequency was abolished in the presence of Cort. **E**. Summary plot of the change in mean normalized sEPSC frequency elicited in CRH neurons by NE in the absence of corticosterone (NE), in the presence of corticosterone (Cort+NE), and in the presence of corticosterone following preincubation of slices in the transcription inhibitor actinomycin D (ActD+Cort+NE). * p < 0.05, ** p < 0.01 compared to baseline sEPSC frequency; #, p < 0.05 compared to NE effect in CRH neurons from unstressed mice.

To test for direct glucocorticoid suppression of NE signaling, we simulated the *in-vivo* stress-induced corticosterone exposure *in vitro* by incubating slices from untreated mice in corticosterone (2 μM) for 30 min prior to returning them to regular aCSF for 2-6 h. Pre-exposure of CRH neurons to corticosterone 2-6 h prior to recordings did not alter basal sEPSC frequency (corticosterone: 1.88 Hz ± 0.32 Hz vs. aCSF control: 1.56 Hz ± 0.10 Hz, n = 16 and 47 respectively, p = 0.70) or amplitude (corticosterone: 19.11 pA ± 1.32 pA vs. aCSF control: 19.13 pA ± 0.50 pA, p = 0.49). However, like acute stress, *in-vitro* corticosterone preincubation suppressed the NE-induced increase in sEPSC frequency 2-6 h later (136.1 ± 11.2% of baseline, n = 11, p < 0.05 compared to vehicle-treated slices) (Fig. 2A, B). These findings suggest that the acute restraint stress-induced desensitization of CRH neurons to NE is due to stress-induced circulating corticosterone.

Corticosterone activates both nuclear glucocorticoid receptors to exert genomic actions and membrane-associated receptors to exert both genomic^20^ and non-genomic^18,21-23^ actions. Increasing evidence suggests that neural plasticity can be rapidly induced non-genomically by membrane glucocorticoid receptors^22-24^. To determine whether the corticosterone suppression of NE-induced excitation of CRH neurons is mediated by a rapid glucocorticoid action, we incubated mouse hypothalamic slices in corticosterone (2 μM) for 5 min prior to co-application of corticosterone and NE (100 μM) for 5 min. Corticosterone application alone caused a significant decrease in sEPSC frequency (88.3% ± 3.2% of baseline, p < 0.05, paired Student’s *t* test, n = 7), which was shown previously to be mediated by retrograde endocannabinoid suppression of glutamate release^22,25,26^. Corticosterone preincubation for 5 min prior to co-application of NE completely blocked the NE-induced increase in sEPSC frequency and resulted in a significant decrease in sEPSC frequency (81.9 ± 16.8% of baseline; p < 0.05, n = 8) (Fig 2C-E). Next, to determine whether the rapid glucocorticoid desensitization of the NE effect is dependent on a transcription-dependent mechanism, we preincubated the slices in the transcription inhibitor actinomycin D (25 μM) for 30 min prior to corticosterone (1 μM) and NE (100 μM) application. Blocking transcription failed to reverse the glucocorticoid suppression of the norepinephrine facilitation of sEPSC frequency (99.37 ± 8.53% of baseline; p = 0.92 compared to NE effect in corticosterone, n = 8) (Fig. 2E). Unexpectedly, inhibiting translation with cycloheximide (100 μM) for 30 min blocked the corticosterone suppression and restored the NE facilitation of sEPSCs (223.10 ± 60.36% of baseline; p = 0.02 compared to NE effect in cort, n = 8) (Fig. S.3), suggesting a dependence on dendritic protein synthesis. The stress desensitization of CRH neurons to NE excitation, therefore, was mediated by a rapid non-genomic corticosteroid signaling mechanism.

To test whether classical glucocorticoid receptors participate in the corticosteroid-induced desensitization of NE excitation of CRH neurons, we used a transgenic mouse model in which the glucocorticoid receptor (GR) is conditionally knocked out in PVN neurons, the CRH-eGFP; sim1GR^-/-^ (GR^-/-^) mouse^27^. Previous studies showed that these mice show an increase in ACTH and corticosterone secretion in response to acute stress due to an impaired glucocorticoid negative feedback^28^, and that PVN CRH neurons in these mice do not respond to rapid glucocorticoid actions^26^. In CRH neurons from loxP-GR control mice lacking cre recombinase (GR^+/+^), corticosterone application (1 μM) alone for 10 min induced a decrease in sEPSC frequency (p < 0.05, N = 8), which was not seen in CRH neurons from GR^-/-^ mice (p = 0.92, n = 8) (Fig. S3B), consistent with our previous findings^22,26^. In CRH neurons from GR^-/-^ mice, corticosterone (1 μM) failed to abolish the NE-induced increase in sEPSC frequency (303.11% ± 20.22% of baseline, p < 0.01, n = 5) (Fig. S3B). Similarly, prior restraint stress (30 min) *in vivo* did not block the NE-induced increase in sEPSC frequency recorded subsequently in CRH neurons from GR^-/-^ mice (208.97% ± 33.88% of baseline, p < 0.01, n = 5) (Fig. S3B). These data, therefore, indicate that depletion of GR prevents the corticosteroid desensitization of CRH neurons to excitation by NE, suggesting that the rapid glucocorticoid desensitization of α1 adrenoreceptors is dependent on the nuclear GR.

### Glucocorticoid desensitization of CRH neurons to NE excitation is mediated by ligand-dependent endocytosis

Corticosterone has been reported to regulate glutamate receptor trafficking in pyramidal neurons^29-31^. Here, we tested whether the rapid glucocorticoid desensitization of CRH neurons to NE is caused by dynamin-dependent α1 receptor endocytosis. Blocking endocytosis with a non-competitive dynamin GTPase inhibitor, Dynasore (80 *µ*M) ^32,33^, applied in the bath for 10 min had no effect on baseline sEPSC frequency (p = 0.875, n = 10), but reversed the corticosterone (2 μM) suppression of the NE (100 μM)-induced increase in sEPSC frequency (259.4% ± 61.3% of baseline, p < 0.05, n = 10) (Fig. 3A-C). This suggested that the rapid glucocorticoid desensitization of the CRH neurons to NE excitation is mediated by a dynamin-dependent internalization of α1-adrenoceptors.

**Fig. 3.**
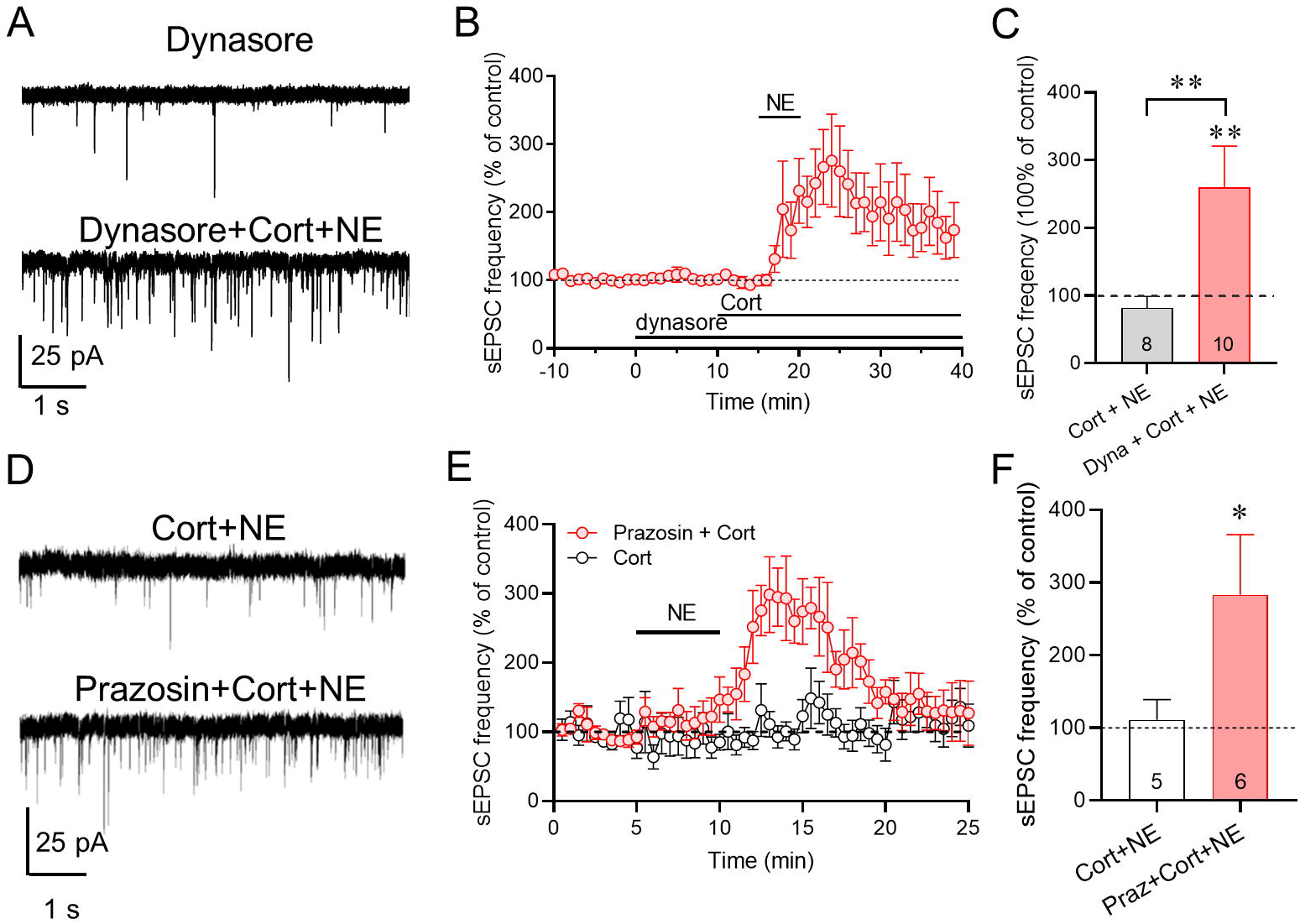
The corticosterone-induced desensitization of CRH neurons to NE excitation is dependent on ligand-mediated endocytosis. **A**. Representative recordings of sEPSCs recorded in CRH neurons at baseline in the dynamin inhibitor Dynasore and after NE application following Dynasore and corticosterone co-application (Dynasore + Cort + NE). **B**. Time course of the change in mean normalized sEPSC frequency showing restoration of the NE-induced increase in sEPSC frequency by Dynasore preapplication prior to Cort and NE application. **C**. Summary plot of NE effect on mean normalized sEPSC frequency in CRH neurons in corticosterone alone (Cort + NE) and in dynasore and corticosterone (Dyna + Cort + NE). Blocking dynamin-dependent endocytosis reversed the corticosterone blockade of the NE-induced increase in sEPSC frequency. **D**. Representative recordings of the sEPSC response to NE 3 h following preincubation of slices in corticosterone (Cort) and in the α1 adrenoreceptor antagonist prazosin and corticosterone (Prazosin + Cort). **E**. Time course of the mean normalized sEPSC frequency showing that the corticosterone-induced desensitization to NE (Cort) was blocked by co-incubation of slices in corticosterone with the α1 adrenoreceptor antagonist prazosin (Prazosin + Cort). **F.** Summary histogram of the increase in sEPSC frequency in response to NE following pretreatment with corticosterone (Cort) or with prazosin and corticosterone (Praz + Cort + NE). The corticosterone suppression of the NE-induced increase in sEPSC frequency was prevented by cotreatment with the α1 adrenoreceptor antagonist prazosin. * p < 0.05, ** p < 0.01.

G protein-coupled receptors undergo ligand-dependent internalization via β-arrestin-triggered endocytosis. α1 adrenoreceptors are tonically activated in PVN CRH neurons by basal NE release (Fig. S. 3C). To determine whether glucocorticoid desensitization of NE signaling in CRH neurons depends on ongoing ligand-mediated α1 receptor internalization, we tested for the dependence of the corticosterone suppression of the NE response on basal adrenoreceptor activation by preincubating slices in the α1 adrenoreceptor antagonist prazosin (10 μM) for 10 min prior to a 5-min coincubation in prazosin and corticosterone (1 μM) for 5 min and 30 min of washout of both. We reasoned that a 30-min washout is sufficient to reverse the prazosin blockade of α1 receptors but not to reverse the glucocorticoid desensitization of the α1 receptors, since a 5-min application of corticosterone desensitizes CRH neurons to NE for over 4 h (unpublished observation). Norepinephrine application (100 μM) following the corticosterone (1 μM) and prazosin (10 μM) co-incubation caused a robust increase in sEPSC frequency (283% +/- 33% of baseline, p < 0.05, n = 6) (Fig. 3D-F). This indicated that blocking α1 receptor activation during corticosterone exposure prevented the glucocorticoid-induced desensitization of the NE response and supports the idea that corticosterone facilitates the ligand-dependent internalization of the α1 adrenoreceptors by endogenous NE.

### Corticosterone facilitates ARα1b internalization

Our electrophysiological data suggest, therefore, that corticosterone facilitates the rapid internalization of α1 adrenoreceptors that leads to the desensitization of the CRH neurons to NE. The dominant form of α1 adrenoreceptor in the CRH neurons is the α1b receptor^13^. To test directly whether the glucocorticoid-induced desensitization to NE is due to α1 adrenoreceptor internalization, we performed a live-cell imaging analysis using the overexpression of a GFP-tagged α1b adrenoreceptor (ARα1b) in an embryonic hypothalamic cell line, MHYPOE N42 (N42) cells, which express both CRH and membrane glucocorticoid receptors^20,34,35^. A 20-min treatment of the cells with corticosterone (2 μM) had no effect on the membrane localization of ARα1b receptors in the N42 cells (Fig. 4A, B), which confirmed that corticosterone alone does not induce α1 adrenoreceptor internalization. We next tested whether NE causes ligand-dependent ARα1b internalization and if corticosterone facilitates the ARα1b internalization. Norepinephrine alone (1 μM) caused a decrease in the membrane localization of the ARα1b-GFP, indicating internalization of membrane α1 receptors (Fig. 4A, B). Co-application of NE and corticosterone (2 μM) caused a ∼2-fold decrease in the ARα1b membrane localization compared to NE alone (Fig. 4A, B). These data indicate that corticosterone does not directly induce α1 adrenoreceptor internalization, but facilitates the internalization of the receptors induced by NE.

**Figure 4.**
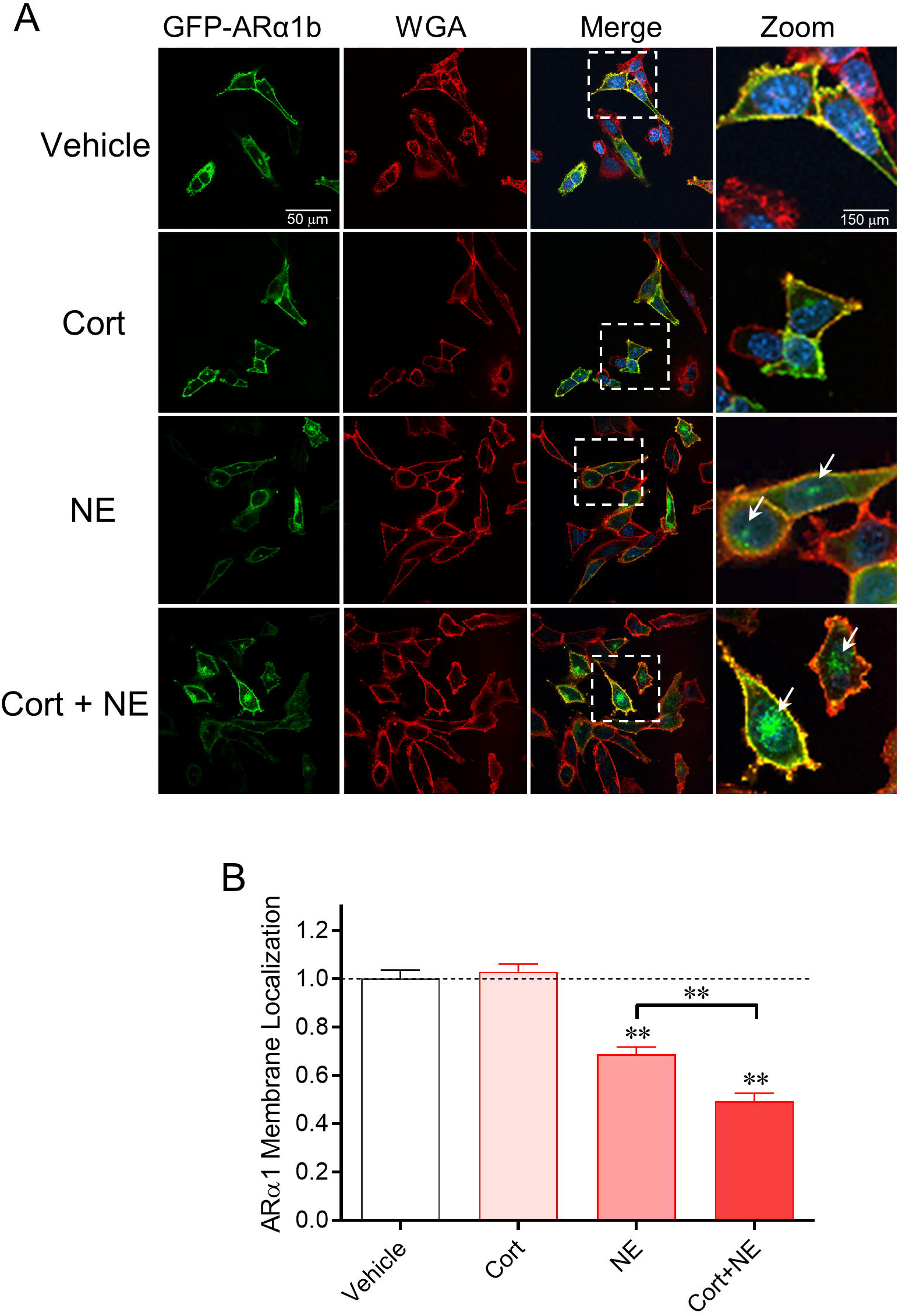
Corticosterone facilitates ligand-mediated α1b adrenoreceptor internalization. **A.** Serum-stripped N42 hypothalamic cells were transiently transfected with ARα1b-eGFP (green) and co-stained with the membrane marker wheatgerm agglutinin (WGA-594, red) and the chromatin marker DAPI (blue). **Vehicle**: Expression of ARα1b is localized largely in the plasma membrane under baseline conditions. **Cort**: ARα1b localization is maintained mostly in the membrane following corticosterone treatment. **NE**: NE treatment caused an increase in ARα1b localization inside the cells, seen as an accumulation of GFP staining in intracellular clusters (“hot spots”, white arrows) and a decrease in membrane co-localization with WGA-594. **Cort+NE**: Co-application of corticosterone with NE increased the ARα1b internalization, as seen by the decrease in ARα1b and WGA-594 membrane co-localization and the increase in the size of the ARα1b hot spots in the cytoplasm (white arrows). **B**. Summary plot of the normalized ARα1b membrane localization in N42 cells in vehicle and after incubation in corticosterone alone (Cort), NE alone (NE), and corticosterone and NE (Cort + NE). Co-localization of ARα1b with WGA was calculated as a Pearson’s coefficient comparing pixel intensity normalized to vehicle. ** p < 0.01.

### Glucocorticoid suppression of NE excitatory drive is specific to physiological stress activation

Previous studies have reported that noradrenergic afferents to the PVN mediate the physiological stress-induced excitation of the PVN CRH neurons, but that this input is not required for CRH neuron activation by psychological stress^36^. Using an immune challenge as a physiological stressor and a predator odor as a psychological stressor, we tested whether a previous stress exposure (restraint) would have a differential effect on HPA activation by a physiological stress compared to a psychological stress (Fig. 5A). Mice that received no prior restraint stress and only LPS injection (100 mg/kg I.P.) or predator odor exposure (30 min) showed an increase in blood corticosterone compared to non-stressed, I.P. saline-injected mice (Fig. 5B). Prior exposure to restraint stress caused a significant suppression of the LPS-induced increase in corticosterone, but had no effect on the corticosterone response to predator odor stress (Fig. 5B). Non-stressed, handled mouse corticosterone levels were used as baseline levels (Fig. 5B).

**Figure 5.**
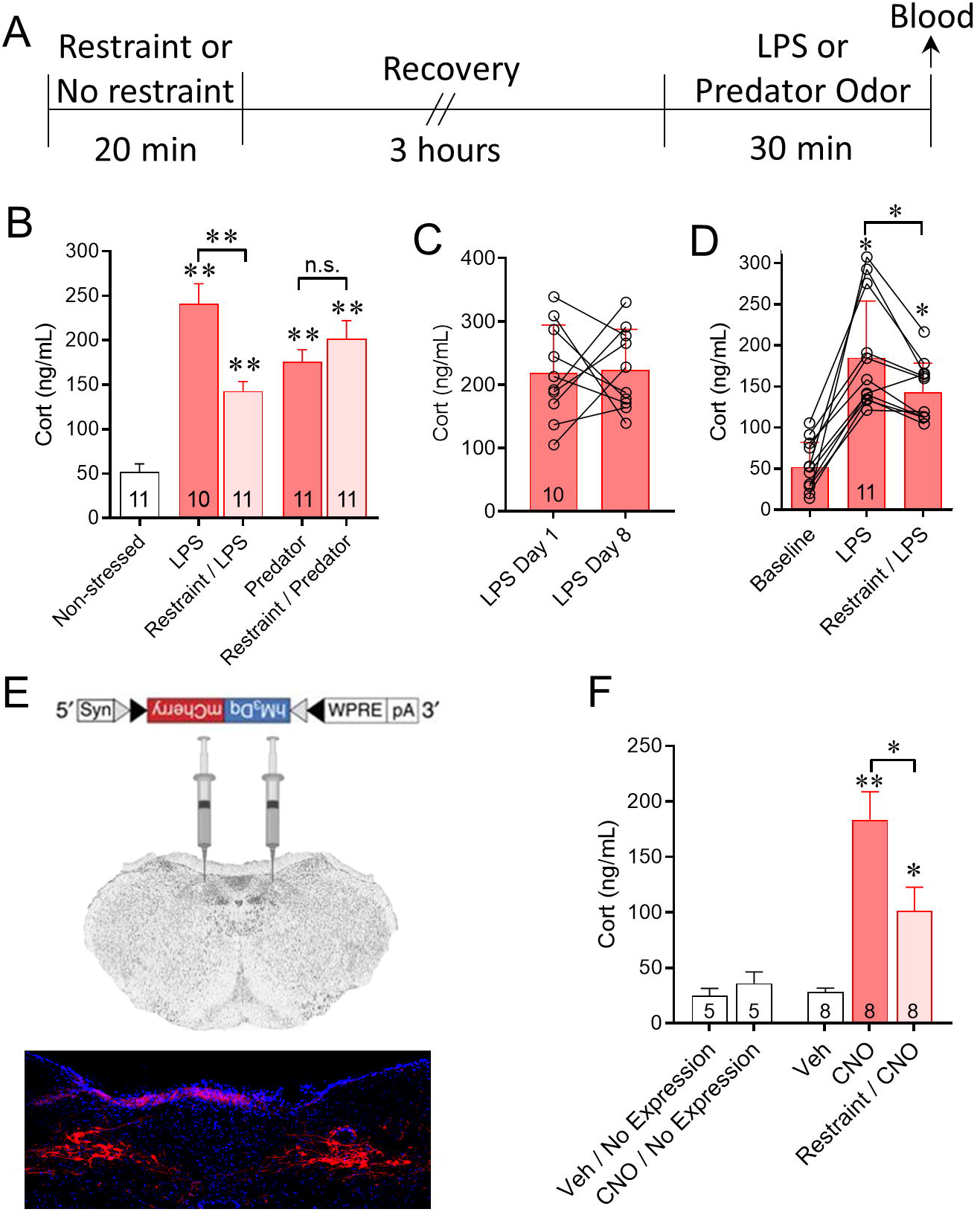
Prior stress exposure differentially suppresses the HPA response in a stress modality-specific manner via desensitization to noradrenergic activation. **A**. Experimental protocol used for sequential stress exposures shown in B. An initial restraint stress or home cage control (No restraint) was followed 3 h later by either I.P. LPS injection or predator odor exposure for 30 min, after which blood was taken. **B**. The LPS-induced increase in corticosterone (LPS) compared to vehicle-injected, unstressed controls (Non-stressed) was suppressed when preceded by a restraint stress (Restraint/LPS). The predator odor-induced increase in corticosterone (Predator) was unaffected by prior restraint stress (Restraint/Predator). (ANOVA: p<0.0001, Tukey’s post-hoc: **, p < 0.01. **C**. The average LPS-induced increase in plasma corticosterone was unaffected by a previous LPS treatment 7 days earlier (LPS Day 1 vs LPS day 8, p = 0.88 n = 10), which excluded within-subject order effects with sequential injections at a 1-week interval. **D**. A cohort of animals had plasma corticosterone levels measured using a within-subject analysis after I.P. saline injection (Baseline), followed 3 days later with I.P. LPS injection 3 h after a 30-min restraint (restraint/LPS), followed 7 days later with LPS injection (LPS) alone. LPS injection following restraint caused a significantly smaller increase in plasma corticosterone that LPS injection without prior restraint. **E**. Cre-dependent excitatory DREADD ((hM_3_Dq)/mCherry)-expressing AAV9 injected bilaterally into the NTS of TH-Cre mice induced mCherry expression in A2 noradrenergic neurons after 3 weeks. **F**. In mice in which DREADD expression was detected in the NTS in *post-hoc* histological controls, CNO injection stimulated an HPA corticosterone response (CNO) compared to vehicle injection (Veh), which was suppressed by prior restraint stress (Restraint/CNO). Vehicle injections and CNO injections in mice in which no virus was detected in *post-hoc* brain sections (Veh/No Expression and CNO/No Expression) failed to elicit an increase in corticosterone. ** p < 0.01, * p < 0.05, Tukey’s multiple comparisons following 1-way ANOVA.

We also tested for an acute stress-induced desensitization of the HPA axis response to LPS injection using a within-subject repeated-measures paradigm to eliminate individual, litter, or cohort variability. First, to test for an order effect of LPS exposure, control mice were stressed with I.P. LPS injections twice in succession, with a seven-day recovery period between injections. No significant difference in blood corticosterone levels between the two responses was found (p=0.89, n=10) (Fig. 5C). To test the effect of acute stress on subsequent LPS-induced corticosterone release, animals were restrained for 30 min, allowed to recover for 3 h, then injected I.P. with LPS (100 *µ*g/kg). The same animals were then injected seven days later with LPS without prior restraint. Significantly lower corticosterone levels were seen in response to LPS injection following restraint than in the same animals injected with LPS only (p < 0.05, n = 11) (Fig. 5D).

The LPS-induced stress response activates noradrenergic inputs from the NTS^36^. We next tested whether the acute stress-induced desensitization of the HPA axis response to LPS is due to desensitization to NTS noradrenergic afferents using a chemogenetic strategy. We expressed an excitatory DREADD in A2 noradrenergic neurons by injecting Cre-dependent AAV9-synhM3Dq-mCherry into the NTS of tyrosine hydroxylase (TH)-Cre mice (Figure 5E). Following a 2-week recovery, mice were subject to a 30-min restraint stress, followed 3 hours later with CNO injection (5 mg/kg, I.P.) and blood collection after another 30 min. Restraint-stressed mice responded to CNO with significantly lower levels of corticosterone compared to stress-naïve mice (Figure 5F, p < 0.05, n = 8). The stress-induced suppression of the corticosterone response to NTS noradrenergic neuron activation was comparable to the stress-induced suppression of the HPA response to LPS. These data suggest that acute stress causes specific desensitization of the norepinephrine-induced activation of the HPA axis by physiological, but not psychological stress.

## DISCUSSION

There is considerable evidence that indicates that noradrenergic inputs to the PVN are instrumental in the activation of the HPA axis^3,37^ via α1-adrenoceptor excitation of the CRH neurons^8,38,39^. We have shown that this noradrenergic activation of PVN CRH neurons is mediated by a retrograde neuronal-glial signaling mechanism that results in stimulation of local excitatory and inhibitory synaptic circuits^10^. Increased circulating corticosterone modulates α1 adrenoceptor-mediated neuronal excitability^15,40^ and α1 adrenoceptor mRNA expression in the hypothalamus and hippocampus^13,41^. Here, we show that acute stress-induced glucocorticoid feedback rapidly desensitizes CRH neurons to NE excitation by facilitating ligand-dependent α1 adrenoreceptor internalization, and that this specifically suppresses the HPA response to physiological, but not psychological, stress.

Norepinephrine causes a robust activation of glutamatergic synaptic inputs and a less robust activation of GABAergic synaptic inputs to PVN CRH neurons via an α1 adrenoreceptor-dependent retrograde neuronal-glial signaling mechanism, resulting in robust excitation of the CRH neurons^10^. Our findings here indicate that pre-exposure of the mouse to acute stress or preincubation of brain slices in corticosterone suppresses the NE activation of glutamate and GABA circuits by facilitating α1 adrenoreceptor internalization, which effectively desensitizes the CRH neurons to NE. We propose a model of glucocorticoid-induced desensitization of α1 adrenoreceptors via receptor internalization that is supported by the following findings: 1) the NE response in CRH neurons is restored by inhibiting glucocorticoid synthesis during the acute stress *in vivo*; 2) the NE response is preserved by inhibiting endocytosis during corticosterone exposure *in vitro*; and 3) corticosterone facilitates the ligand-induced internalization of the α1b receptor. That corticosterone by itself did not induce changes in membrane receptor trafficking in cultured cells and that the NE desensitization by corticosterone was blocked by blocking α1 receptors during corticosterone exposure in brain slices suggest that corticosterone *per se* does not induce α1 receptor internalization, but rather it regulates α1 receptor trafficking following ligand-induced internalization. The CRH neurons were almost entirely desensitized to bath-applied NE with even 5 min of corticosterone exposure, which suggests that the membrane α1 adrenoreceptor undergoes nearly complete internalization and recycling within 5 min, and that corticosterone acts rapidly to prevent α1 receptor trafficking back to the membrane once it has been internalized.

The rapid regulation of α1 adrenoreceptor trafficking by corticosterone suggests a non-genomic mechanism of action of the steroid, which was supported by the ineffectiveness of the transcription blocker actinomycin D to reverse the corticosterone effect. Nevertheless, the effect of the steroid was lost in CRH neurons from GR knockout mice, which suggests that it is dependent on the nuclear GR. We have reported a rapid glucocorticoid-induced endocannabinoid suppression of excitatory synapses in PVN CRH neurons and magnocellular neurons that is also dependent on the GR, but does not require gene transcription and is mediated by an intracellular signaling cascade^22,25,26,42^. The rapid glucocorticoid suppression of excitatory synaptic inputs was also observed here, and was also reversed in CRH neurons from GR knockout mice. It is reasonable to envision the membrane transduction mechanism responsible for the rapid glucocorticoid regulation of α1 receptor trafficking in CRH neurons as the same as or as a branch of the membrane glucocorticoid receptor signaling mechanism that leads to endocannabinoid synthesis at glutamate synapses^42^ and ligand-independent nuclear translocation of the GR^20^, although further experiments are required to test this. An unexpected finding here was that the corticosterone suppression of the NE excitatory effect was reversed by inhibiting translation with cycloheximide (Fig. S3), since this suggests that a component of the glucocorticoid regulation of α1 receptor trafficking may depend on constitutive protein synthesis.

Corticosterone facilitates glutamate neurotransmission by regulating AMPA receptor trafficking in pyramidal neurons of the hippocampus and prefrontal cortex^29-31^. In addition, corticosterone slowly suppressed β adrenoreceptor-induced enhancement of AMPA currents^43^, and diminished the β adrenoreceptor-induced facilitation of synaptic strength in the basolateral amygdala^44^ and hippocampal dentate gyrus^45^. Serum- and glucocorticoid-inducible kinase (SGK), guanosine nucleotide dissociation inhibitor (GSI), and Rab family small-molecule G proteins were reported to mediate the corticosterone regulation of AMPA receptor trafficking in the prefrontal cortex^31,46^. In hippocampal cell cultures, corticosterone stimulated the tyrosine kinases Pyk2 and Src to cause a rapid increase in PSD95 clustering and NMDA receptor surface expression by activating a membrane receptor and downstream signaling^47^. It is reasonable to envision that corticosterone also facilitates α1-adrenoceptor internalization by triggering intracellular kinase cascades and/or destabilizing scaffolding proteins, although this remains to be determined.

The three subtypes of α1 adrenoceptors, α1a, α1b, and α1d receptors, are all expressed in the PVN. However, the α1b is most likely responsible for mediating the NE response because 1) α1a mRNA expression is low in the PVN^48^ and is not regulated by glucocorticoids^13^; 2) only 37% of CRH neurons express α1d adrenoceptor mRNA^49^; and 3) nearly all CRH neurons express α1b adrenoceptor mRNA and its expression is inversely correlated with plasma corticosterone^13^. Native α1b adrenoreceptors are more subject to ligand-induced internalization than α1a receptors. We found that, unlike the α1b adrenoreceptors, α1a receptors expressed in N42 cells were resistant to internalization (Weiss and Tasker, unpublished observation) and corticosterone does not desensitize α1a receptor signaling in the basolateral amygdala (Fu and Tasker, unpublished observation).

Our data suggest that the glucocorticoid desensitization to noradrenergic activation of CRH neurons attenuates HPA activation by physiological stress, but not by psychological stress. This is consistent with systemic stress signals being transmitted to the PVN via ascending noradrenergic afferents from the NTS, whereas psychological stress signals of limbic origin are not thought to transit through the brainstem noradrenergic cell groups to reach the PVN^2-4,13,50^. To our knowledge, this is the first demonstration of a stress modality-specific feedback inhibition of the HPA axis, which takes systemic stress signaling to the HPA axis largely offline and leaves psychological stress activation of the HPA axis intact. The physiological stress activation of the HPA axis is not entirely abolished, with about 50% of the response remaining following stress inactivation. We reported previously the co-release of glutamate with norepinephrine from NTS-derived noradrenergic afferents to CRH neurons^10^. The residual, stress-insensitive component to the response is likely to be mediated by these ascending monosynaptic glutamatergic inputs. The question remains as to the functional significance of the selective suppression of the physiological stress response and sparing of the psychological stress response by prior stress exposure. Are ascending and descending modalities of stress signaling in competition for influence over the stress response? Ascending noradrenergic afferents carry signals generated by systemic stress stimuli, such as immune system, glycemic, and blood pressure signals, whereas descending limbic afferents to the HPA axis carry emotional and cognitive signals from the amygdala, hippocampus, and prefrontal cortex. The ascending systemic afferents, therefore, signal long-lasting physiological states of the organism. It is likely that desensitization of the noradrenergic afferents prevents the tonic activation of the HPA axis by systemic stress signals, protecting the organism from the damaging effects of sustained glucocorticoid exposure, while allowing the phasic activation of the HPA axis by descending limbic signals. The animal is thus able to continue to mount normal transient HPA responses to anticipated or programmed threats to its survival while limiting its exposure to sustained elevated glucocorticoids induced by homeostatic perturbation. Further studies are required to determine the physiological significance of the stress modality-specific regulation of the HPA axis in the context of susceptibility/resilience to systemic stressors such as immune challenge.

## ACKNOWLEDGMENTS

This study was supported by NIH grants 2R01 MH066958 and R01 MH119283, the Catherine and Hunter Pierson Chair in Neuroscience, and the Tulane Undergraduate Research in Neuroscience Program.

## METHODS

### Animals

We used 6-10-week-old male mice for these experiments; all animals and procedures were approved by the Tulane Institutional Animal Care and Use Committee (IACUC) according to National Institutes of Health (NIH) guidelines. All mice were housed under controlled temperature and humidity on a 12:12 light/dark cycle and received food and water *ad libitum*.

Two strains of mice were used for this study, CRH-eGFP transgenic mice and sim1-cre::loxP-GR::CRH-eGFP mice. CRH-eGFP mice were bred on a C57BL/6 background from BAC transgenic mice expressing enhanced green fluorescent protein (eGFP) controlled by the CRH promoter (CRH-eGFP mice). Sim1-cre/loxP-GR::CRH-eGFP mice were bred by crossing the CRH-eGFP mice with conditional glucocorticoid receptor (GR) knockout mice (see Supplemental Methods).

### Acute stress paradigms

#### Restraint stress

Mice were immobilized for 30 min between 9 AM and 10 AM by placing them in a transparent, pliable tapered plastic cone open at the tip to allow for breathing (DecapiCones, Braintree Scientific). Immediately after the 30 min restraint stress, mice were decapitated in the plastic cone without anesthetic and brain slices were prepared.

#### Predator odor stress

Mice were placed in a clean cage between 9 AM and 10 AM and a sponge soaked with bobcat (Lynx rufus) urine (Maine Outdoor Solutions) was placed in a dish in the cage’s water bottle holder. Mice had no direct access to or contact with the sponge. Following a 30-min exposure to the bobcat urine, they were then placed immediately into a decapicone and decapitated without anesthetic for brain slice preparation.

#### Serial stressors

Mice were first immobilized for 30 min between 9 AM and 10 AM and then returned to their home cage for 3 hours of recovery. They were then exposed to either predator odor for 30 min (see above) or they were injected I.P. with 100 μg/kg lipopolysaccharide (LPS) 45 min prior to blood collection for corticosterone measurement.

### Brain slice preparation and electrophysiology

#### Acute brain slice preparation

Electrophysiological experiments were conducted in acutely prepared hypothalamic slices from 6-9-week-old male CRH-eGFP or sim1-cre::loxP-GR::CRH-eGFP mice. Mice were gently removed from their home cage to a transfer cage and transported to an adjacent room, where they were immobilized in a DecapiCone (Braintree Scientific) and decapitated without anesthesia. Following decapitation, the brain was quickly removed from the skull and cooled in oxygenated, ice-cold artificial cerebrospinal fluid (aCSF) containing (in mM): 140 NaCl, 3 KCl, 1.3 MgSO_4_, 11 Glucose, 5 HEPES, 1.4 NaH2PO_4_, 3.25 NaOH, and 2.4 CaCl_2_, with a pH of 7.2-7.4 and an osmolarity of 290-300 mOsm. Coronal slices (300 μm) containing the PVN were sectioned in cooled aCSF and then maintained in oxygenated aCSF at room temperature for at least 1 h prior to recordings.

#### Electrophysiological recordings

eGFP-expressing CRH neurons in the PVN were targeted for whole-cell recordings using patch pipettes filled with an internal patch solution containing (in mM): 120 potassium gluconate, 10 KCl, 1 NaCl, 1 MgCl_2_, 0.1 CaCl_2_, 5.5 EGTA, 10 HEPES, 2 Mg-ATP, 0.3 Na-GTP. Loose-seal, cell-attached patch clamp recordings were performed with low-resistance pipettes filled with aCSF, as previously described. A high-potassium extracellular solution was used to depolarize neurons and stimulate spontaneous spiking activity, which consisted of (in mM): 133 NaCl, 10 KCl, 1.3 MgSO_4_, 1.4 NaH_2_PO_4_, 2.4 CaCl_2_, 11 glucose, and 5 HEPES; pH was adjusted to 7.2–7.4 with NaOH.

### Receptor trafficking analysis

An immortalized hypothalamic cell line, mHypoE-N42 cells (Cedar Lane, Cellutions Biosystems)^20^, expressing pEGFP-ARα1b was used to visualize intracellular α1b adrenoreceptor trafficking. The cells were treated with 1 *µ*M NE, 2 *µ*M corticosterone:HBC, or both (Sigma Aldrich) for 20 min, and then fixed immediately with 4% paraformaldehyde. They were then stained with the plasma membrane marker WGA Alexa Fluor 594 (ThermoFisher) for 30 s and imaged under confocal microscopy (Nikon A1+, Nikon Instruments, Inc). The Pearson’s coefficient was calculated using Cell Profiler (Broad Institute, Cambridge, MA).

### Statistical analyses

Data are presented as the mean ± standard error of the mean. Excitatory postsynaptic currents (EPSCs) were analyzed for changes in frequency, amplitude and decay time. Statistical significance was determined with the two-tailed, paired Student’s *t*-test for within-cell drug effects and the two-tailed, unpaired Student’s *t*-test for between-group drug effects. A one-way analysis of variance (ANOVA) with *post-hoc* Tukey’s tests were used for comparing corticosterone levels with multiple-group comparisons. Animal numbers are designated as “N” and neuron numbers as “n”.

