## Supplemental Information for "Acute Stress Desensitizes Hypothalamic CRH Neurons to Norepinephrine and Physiological Stress"

**Supplemental Methods**

*Mouse models* - Three mouse models were used in this study, a CRH-eGFP mouse, a triple transgenic sim1-cre::loxP-GR::CRH-eGFP mouse, and a tryrosine hydroxylase (TH)-cre mouse that expresses cre recombinase in catecholaminergic neurons. CRH-eGFP mice were obtained from the Mutant Mouse Regional Resource Center (MMRRC) (stock: Tg(Crh-eGFP)HS57Gsat/Mm, RRID:MMRRC_017058-UCD) at the University of California at Davis; they were used to visualize individual CRH neurons to target for patch-clamp recordings. Transgenic mice were genotyped following the supplier’s protocol with the primers 5’-CTGTCTTGTCGTGGGTGTCCGAT-3’ and 5’-TAGCGGCTGAAGCACTGCA-3’, to produce a 400 bp fragment.

A previous study reported that a CRH-eGFP BAC transgenic mouse from the MMRRC, which is a different strain from the one we used here and previously, expresses eGFP ectopically ^1^. We validated that eGFP is expressed in PVN CRH neurons, and not ectopically, in our mouse line by immunofluorescence in a previous study (Chen et al., 2019). Additionally, here we compared qualitatively the PVN CRH expression in the CRH-eGFP BAC transgenic mouse with the PVN CRH expression in a knock-in CRH-ires-Cre mouse expressing a tdTomato reporter. A cross of the CRH-eGFP mouse and the CRH-ires-Cre::tdTomato mouse (N = 2) showed near-complete overlap of eGFP and tdTomato expression in the PVN (Fig. S4). We preferred to use the CRH-eGFP mouse for our studies because the transgene expression in this mouse is weaker than in the CRH-ires-cre knock-in mouse and less likely, therefore, to interfere with native cell function.

Mice with a conditional knockout of GR in the PVN were generated by crossing a sim1-cre BAC transgenic mouse (Sim1-Cre, kindly provided by Dr Bradford Lowell, Beth Israel Deaconess Medical Center, Boston, MA)^2^ with a loxP-GR transgenic mouse (kindly provided by Dr. Louis Muglia, University of Cincinnati Children’s Hospital, Cincinnati, OH)^3-5^. Sim1-cre:loxP-GR mice show a deletion of GR in the PVN^3,5^. We crossed the sim1-cre/loxP-GR mouse with the CRH-eGFP mouse to produce a sim1-cre::loxP-GR::CRH-eGFP mouse in order to visualize CRH neurons in our brain slices. The pups were genotyped at 2-3 weeks using the primer set: 5’-AATCAGAATTGCTCACTCACAA-3’ (GR 6452–73), 5’ -CAGTGTTACTACTTCCAGTTC-3’ (GR 6670 –50), 5’-TGCTATACGAAGT TATCAGTAC-3’ (LoxPForward), 5’-AAGTGCCTTCTCTACACCTG-3’ (Cre 1123–1004), and 5’-TGCTTATAACACCCTGTTACG-3’ (Cre 982–1002).

TH-cre mouse breeders were purchased from Jackson Laboratory and bred in-house. They were genotyped at 2-3 weeks using the supplier’s suggested primer set.

*Brain slice preparation* - Mice were immobilized in a DecapiCone (Braintree Scientific) and decapitated for brain slice preparation using a rodent guillotine. They were decapitated rapidly, within less than 2 min of removal from their home cage, to avoid stress-elevated circulating corticosterone levels reaching the brain. Mice were killed without anesthesia because anesthetics activate the HPA axis and increase circulating ACTH and corticosterone^6^. Blood corticosterone levels remain low in our hands for >3 min from the start of handling.

Following decapitation, the brain was immersed for ~1 min in oxygenated, ice-cold artificial cerebrospinal fluid (aCSF). The base of the brain was then blocked and the caudal face of the block was glued to the chuck of a vibratome (Leica or Vibratome). Two or three 300-μm coronal slices containing the PVN were sectioned in cooled aCSF and the slices were bisected down the midline and transferred to an incubation chamber. They were maintained in oxygenated aCSF at room temperature for at least 1 h prior to beginning recording sessions to allow for recovery.

*Whole-cell and loose-seal patch clamp recordings –* Hemi-slices were transferred one-at-a-time from the incubation chamber to the recording chamber, where they were submerged and perfused with aCSF at a rate of ~2 ml/min. eGFP-expressing CRH neurons in the PVN were located under fluorescence illumination and subsequently targeted for whole-cell patch clamp recordings using infrared light and differential interference contrast optics at room temperature on a fixed-stage, upright microscope (Olympus BXW51) equipped with a long working distance, water-immersion 40x objective. Patch pipettes with a resistance of 3-6 MΩ were fabricated from borosilicate glass (ID 1.2 mm, OD 1.65 mm; Garner Glass) on a horizontal puller (P-97, Sutter Instr.) and filled with an internal patch solution containing (in mM): 120 potassium gluconate, 10 KCl, 1 NaCl, 1 MgCl_2_, 0.1 CaCl_2_, 5.5 EGTA, 10 HEPES, 2 Mg-ATP, 0.3 Na-GTP; the pH was adjusted to 7.3 with KOH and the osmolarity was adjusted to 300 mOsm with D-sorbitol.

For loose-seal, cell-attached patch clamp recordings, glass pipettes with a resistance of 1-2 MΩ filled with aCSF were used. Cells were recorded at resting potential with minimal current transfer and no exchange of intracellular ions^7,8^. An extracellular aCSF with high [K^+^] (10 mM) was used to stimulate spontaneous spiking activity (in mM): 133 NaCl, 10 KCl, 1.3 MgSO_4_, 1.4 NaH_2_PO_4_, 2.4 CaCl_2_, 11 glucose, and 5 HEPES; pH was adjusted to 7.2–7.4 with NaOH.

*Cell imaging* - Immortalized embryonic mouse hypothalamic cells that express CRH and glucocorticoid receptors (mHypoE-N42, Cedar Lane, Cellutions Biosystems)^9,10^ were transiently transfected with pEGFP-ARα1b (generous gift from Dr. Chris Hague, University of Washington, Seattle, WA) using the Neon® Transfection System (Life Technologies) with 2 pulses, 10 ms, 1600 mV. Cells were placed on coverslips and the media was replaced with media containing charcoal-stripped BSA to eliminate basal steroid content. The cells were treated with 1 µM NE, 2 µM corticosterone:HBC, or both (Sigma Aldrich) for 20 min. Cells were then fixed immediately with 4% paraformaldehyde before staining for 30 s with WGA Alexa Fluor 594 (ThermoFisher) to produce a plasma membrane-specific label. Image analysis was performed on a Nikon A1+ laser confocal microscope (Nikon Instruments, Inc) with a 28 μm pinhole to capture an intracellular focal plane. The average Pearson’s coefficient was calculated in each transfected cell using Cell Profiler (Broad Institute, Cambridge, MA).

*Plasma corticosterone assay* – To measure serum corticosterone concentration, blood (~200 μl) was collected by decapitation (for terminal corticosterone measurement) or by submandibular vein puncture (for serial corticosterone measurement) The blood was allowed to coagulate at room temperature for 90 min and was centrifuged at 2000 × g for 15 min. Serum was collected and samples were stored at -20^o^C until they were shipped to the University of Virginia Center for Research in Reproduction Ligand Assay and Analysis Core (terminal experiments), where corticosterone levels were measured by ^125^I corticosterone radioimmunoassay, or until they were assayed in-house using a corticosterone ELISA (Enzo Life Sciences, Farmingdale, NY) according to the manufacturer’s instructions (serial measurement experiments).

### *Drugs used* – The following drugs were kept at -20^o^C as stock solutions and dissolved in aCSF to their final concentrations on the day of experiments: L-(−)-norepinephrine (+)-bitartrate salt monohydrate (NE, 1 µM, 100 µM, Sigma-Aldrich), picrotoxin (PTX, 50 µM, Tocris), corticosterone (2 µM, Tocris), 3-hydroxynaphthalene-2-carboxylic acid (3,4-dihydroxybenzylidene) hydrazide (Dynasore, 80 µM, Tocris). 2-Methyl-1,2-di-3-pyridinyl-1-propa­none (metyrapone, Tocris) was dissolved in sterile saline, and injected intraperitoneally at a dose of 100 mg/kg body weight.

### *Statistical analyses –* Data are presented as the mean ± standard error of the mean. EPSCs were selected and analyzed with Minianalysis 6.0 (Synaptosoft Inc.). EPSC frequency, amplitude and decay time were compared using the two-tailed paired Student’s *t*-test for within-cell drug effects and the two-tailed, unpaired Student’s *t*-test was used for between-group effects (Prism 7 and Prism 8, Graphpad). A one-way analysis of variance (ANOVA) combined with *post-hoc* Tukey’s tests (SigmaPlot 11.0, Systat Software, Inc.) were used for multiple comparisons.

**Supplemental Figure Legends**

**Figure S1.** Norepinephrine stimulates local glutamate and GABA circuits via postsynaptic α1 receptor activation and a retrograde signaling mechanism in CRH neurons. **A**. The NE-induced increase in sEPSC frequency in CRH neurons is blocked by the α1 receptor antagonist prazosin (1 μM) (Prazosin/NE), the voltage-gated Na^+^ channel blocker TTX (1 μM) (TTX/NE), and postsynaptic intracellular infusion of the G protein blocker GDP-βS (1 mM) (GDP-βS/NE). **B**. The NE-induced increase in sIPSC frequency, but not the NE-induced decrease in sIPSC frequency, is blocked by prazosin (Prazosin/NE) and TTX (TTX/NE), and by intracellular infusion of GDP-βS (GDP-βS/NE). These data together indicate a postsynaptic α1 adrenoreceptor-dependent activation of presynaptic glutamate and GABA neurons. Data taken without modification from Chen et al., 2019.

**Figure S2**. Metyrapone blockade of restraint stress-induced corticosterone secretion. Mice subjected to a 30-min restraint displayed a robust increase in plasma corticosterone levels compared to controls (controls: 15.17 ± 5.92 ng/ml; stressed: 434.87 ± 28.02 ng/ml; p < 0.001, one-way ANOVA, N = 7 and 6, respectively). Administration of the 11-β-hydroxylase inhibitor metyrapone (I.P., 100 mg/kg) 30 min prior to the start of the restraint stress prevented the stress-induced increase in plasma corticosterone (vehicle: 491.51 ± 21.96 ng/ml; metyrapone: 32.43 ± 3.82 ng/ml); p < 0.05, one-way ANOVA, N = 8 and 7, respectively). ** p < 0.01 compared to unstressed control group.

**Figure S3.** Dependence of corticosterone suppression of NE effect on a transcription-independent, GR-dependent signaling mechanism. **A**. A 5-min application of corticosterone (1 μM) followed immediately with NE application (100 μM) blocked the NE-induced increase in sEPSC frequency (Cort + NE). The corticosterone-induced suppression was not reversed by blocking gene transcription with actinomycin D (25 µM) (ActD + Cort + NE), but was reversed by blocking protein translation with cycloheximide (100 µM) (CHX + Cort + NE). **B**. The corticosterone suppression of the NE facilitation of glutamate release was dependent on the nuclear glucocorticoid receptor. The *in-vitro* corticosterone suppression of the NE-induced increase in sEPSC frequency (Cort/NE-GRKO) and the *in-vivo* stress-induced suppression of the NE response (Stress/NE-GRKO) were reversed in CRH neurons from the conditional GRKO mouse. **C**. Blocking a1 adrenoreceptors with prazosin in the absence of exogenous agonist caused a significant reduction in sEPSC frequency, suggesting a tonic activation of the a1 receptors with endogenous NE. ** p < 0.01, * p < 0.05 compared to baseline sEPSC frequency.

**Figure S4.** CRH reporter transgene expression in two mouse models. A cross between the CRH-eGFP BAC transgenic mouse and the CRH-ires-cre::tdTomato mouse showed nearly complete overlap of the expression of the two fluorescence transgenes, eGFP and tdTomato.

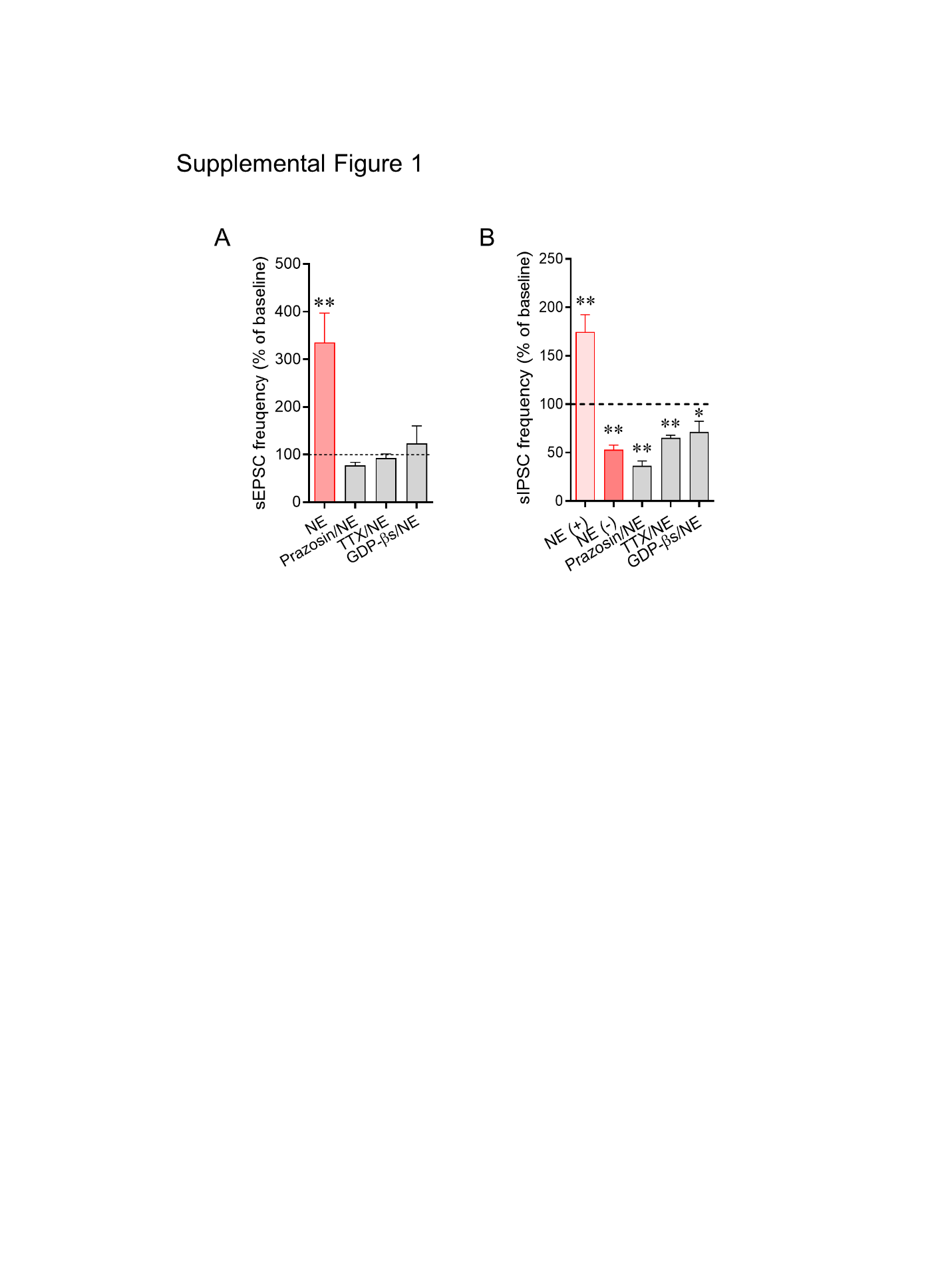

**Figure S1.** Norepinephrine stimulates local glutamate and GABA circuits via postsynaptic α1 receptor activation and a retrograde signaling mechanism in CRH neurons. **A**. The NE-induced increase in sEPSC frequency in CRH neurons is blocked by the α1 receptor antagonist prazosin (1 μM) (Prazosin/NE), the voltage-gated Na^+^ channel blocker TTX (1 μM) (TTX/NE), and postsynaptic intracellular infusion of the G protein blocker GDP-βS (1 mM) (GDP-βS/NE). **B**. The NE-induced increase in sIPSC frequency, but not the NE-induced decrease in sIPSC frequency, is blocked by prazosin (Prazosin/NE) and TTX (TTX/NE), and by intracellular infusion of GDP-βS (GDP-βS/NE). These data together indicate a postsynaptic α1 adrenoreceptor-dependent activation of presynaptic glutamate and GABA neurons. Data taken without modification from Chen et al., 2019.

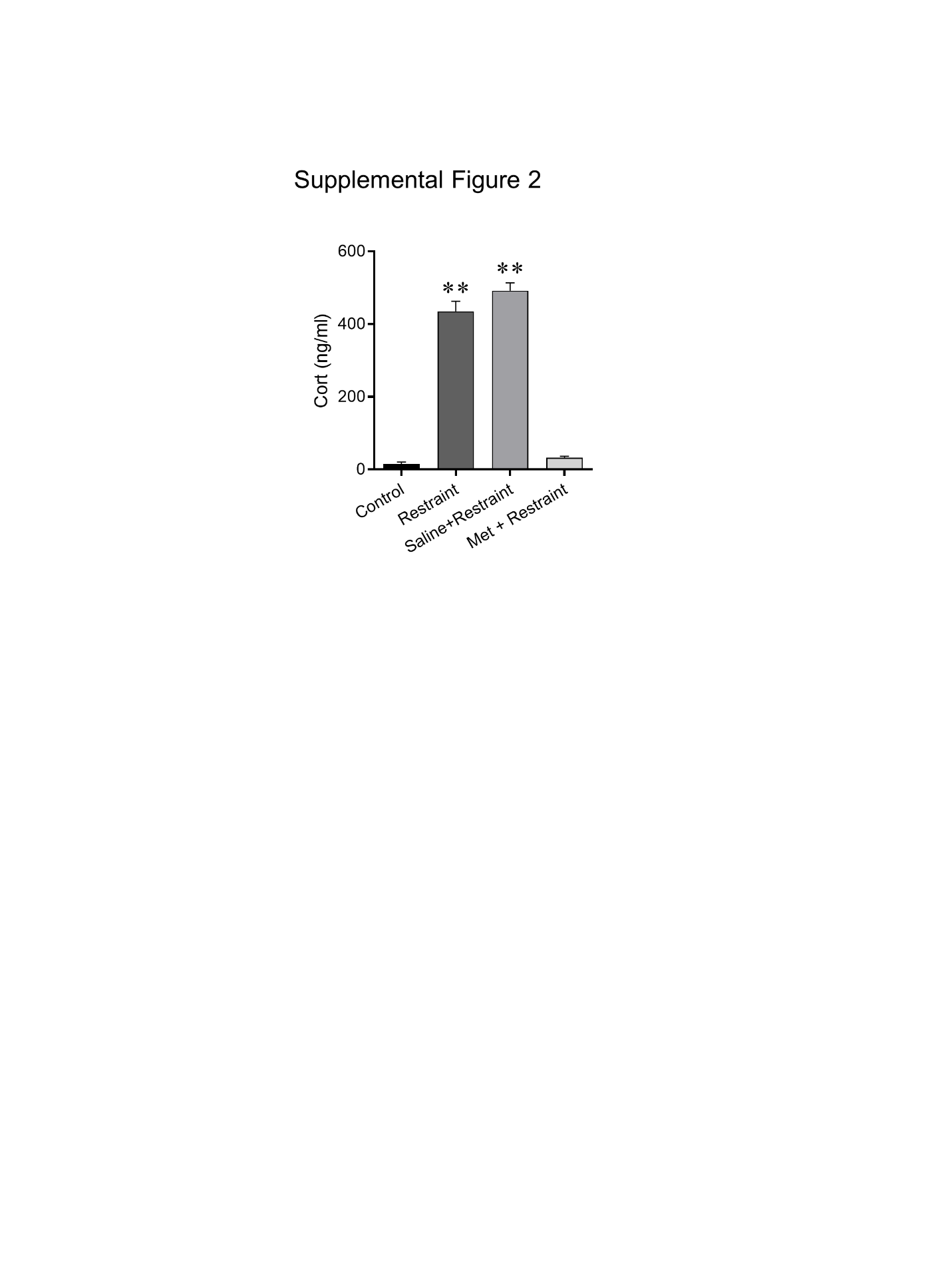

**Figure S2**. Metyrapone blockade of restraint stress-induced corticosterone secretion. Mice subjected to a 30-min restraint displayed a robust increase in plasma corticosterone levels compared to controls (controls: 15.17 ± 5.92 ng/ml; stressed: 434.87 ± 28.02 ng/ml; p < 0.001, one-way ANOVA, N = 7 and 6, respectively). Administration of the 11-β-hydroxylase inhibitor metyrapone (I.P., 100 mg/kg) 30 min prior to the start of the restraint stress prevented the stress-induced increase in plasma corticosterone (vehicle: 491.51 ± 21.96 ng/ml; metyrapone: 32.43 ± 3.82 ng/ml); p < 0.05, one-way ANOVA, N = 8 and 7, respectively). ** p < 0.01 compared to unstressed control group.

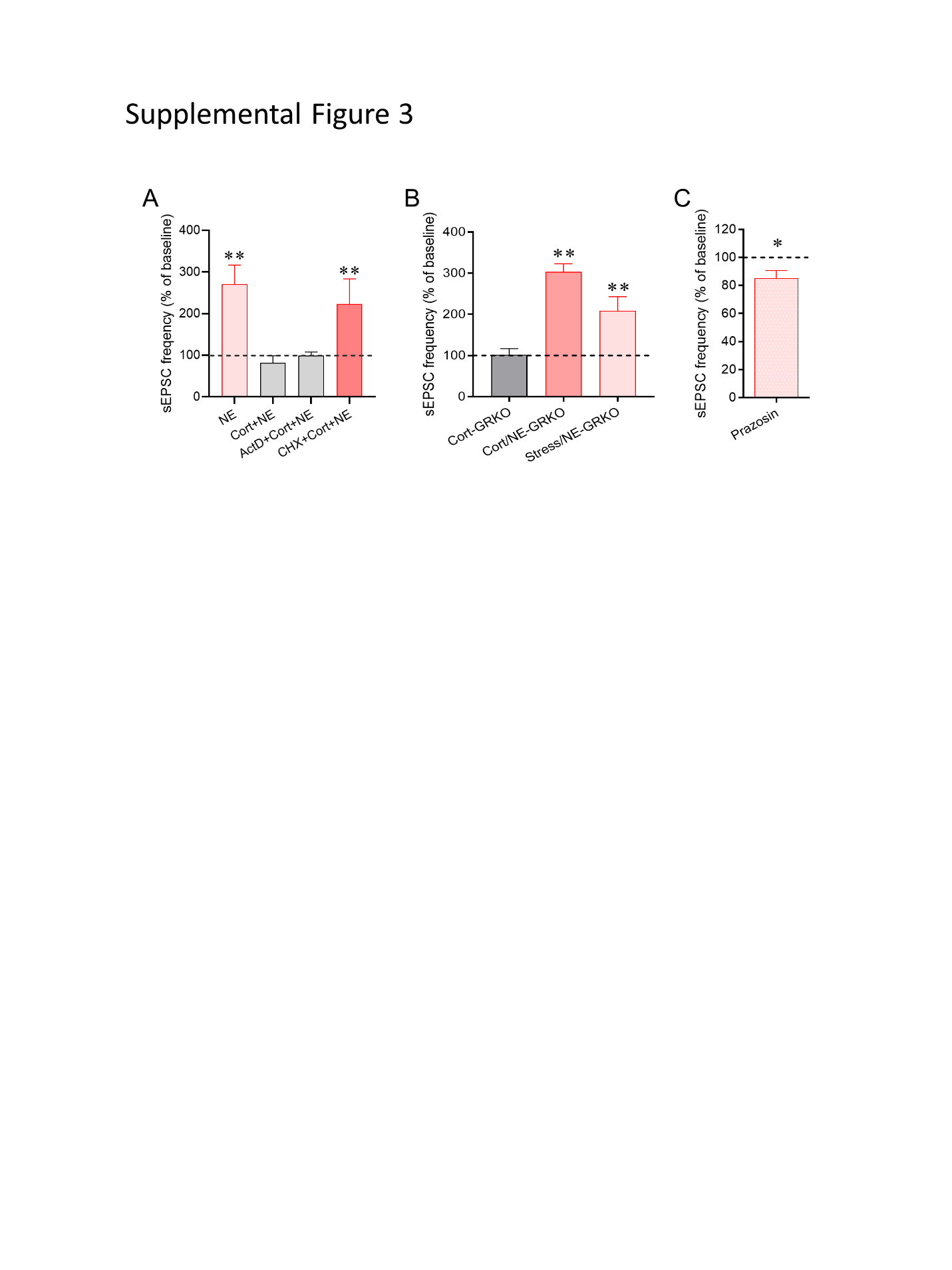

**Figure S3.** Dependence of corticosterone suppression of NE effect on a transcription-independent, GR-dependent signaling mechanism. **A**. A 5-min application of corticosterone (1 μM) followed immediately with NE application (100 μM) blocked the NE-induced increase in sEPSC frequency (Cort + NE). The corticosterone-induced suppression was not reversed by blocking gene transcription with actinomycin D (25 µM) (ActD + Cort + NE), but was reversed by blocking protein translation with cycloheximide (100 µM) (CHX + Cort + NE). **B**. The corticosterone suppression of the NE facilitation of glutamate release was dependent on the nuclear glucocorticoid receptor. The *in-vitro* corticosterone suppression of the NE-induced increase in sEPSC frequency (Cort/NE-GRKO) and the *in-vivo* stress-induced suppression of the NE response (Stress/NE-GRKO) were reversed in CRH neurons from the conditional GRKO mouse. **C**. Blocking a1 adrenoreceptors with prazosin in the absence of exogenous agonist caused a significant reduction in sEPSC frequency, suggesting a tonic activation of the a1 receptors with endogenous NE. ** p < 0.01, * p < 0.05 compared to baseline sEPSC frequency.

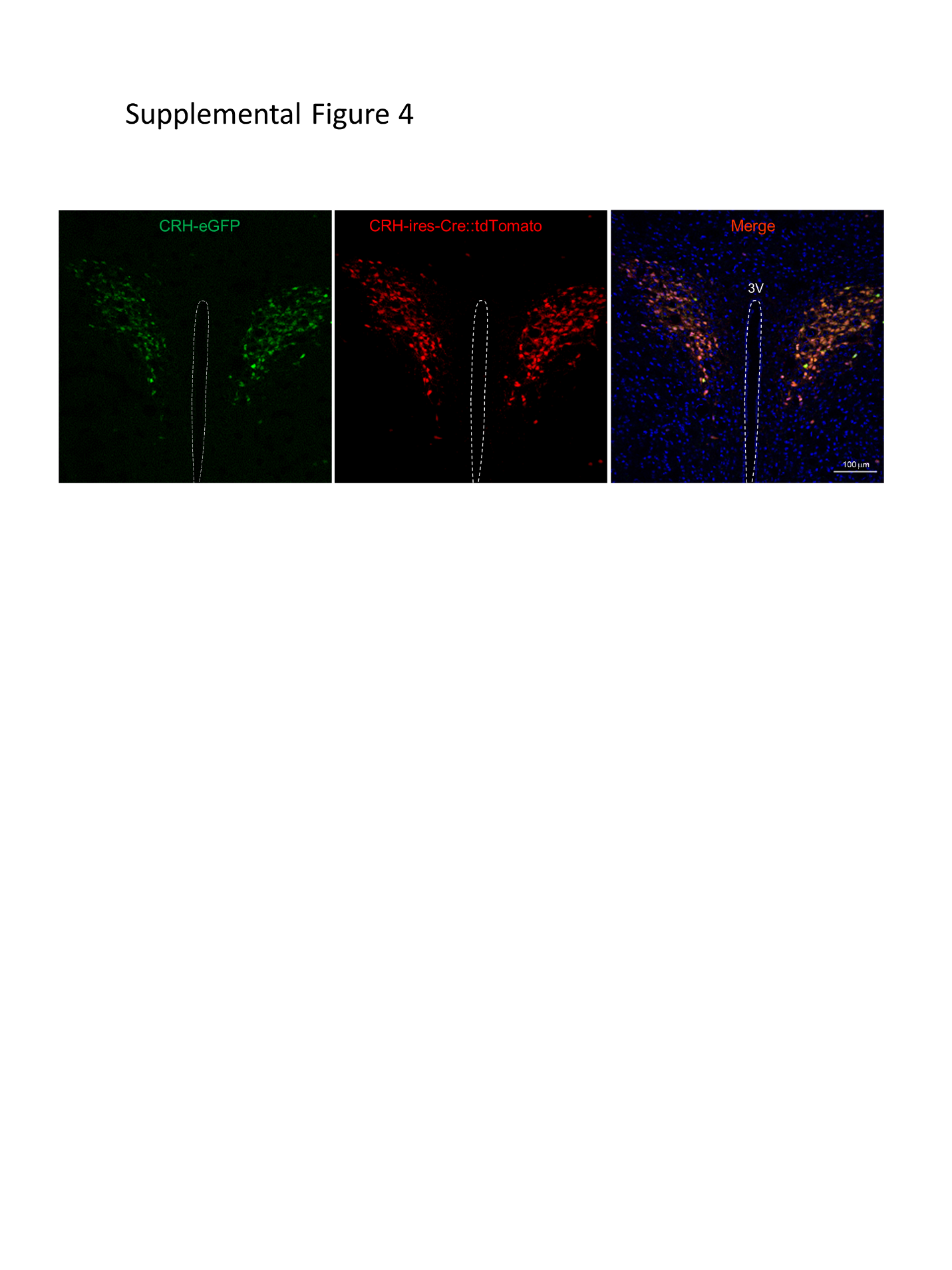

**Figure S4.** CRH reporter transgene expression in two mouse models. A cross between the CRH-eGFP BAC transgenic mouse and the CRH-ires-cre::tdTomato mouse showed nearly complete overlap of the expression of the two fluorescence transgenes, eGFP and tdTomato.
